# Evaluating the Importance of Glucagon in the Insulin-Glucose Regulatory System: A Mechanistic Modeling Approach

**DOI:** 10.1101/2024.07.31.606029

**Authors:** Mackenzie Dalton, Emmanuel Asante-Asamani, James Greene

## Abstract

The dynamics of insulin and glucose are tightly regulated. The pancreatic islets of Langerhans contain both beta and alpha cells which produce insulin and glucagon, respectively. Insulin is the only hormone in the body that lowers blood glucose levels by acting like a key for glucose to enter cells. Without insulin, cells cannot utilize glucose, their primary source of energy. In contrast, glucagon functions as a hormone which elevates blood glucose levels by promoting the breakdown of glycogen in the liver. Maintaining blood glucose within a safe range is vital since both excessively high and low levels can be life-threatening (hyperglycemia and hypoglycemia, respectively), and these two hormones work together to achieve this balance. In this work we aim to underscore the significance of glucagon in the insulin-glucose regulatory system. We construct a three-compartment mechanistic model that includes insulin, glucose, and glucagon, which is then validated by fitting to publicly available from an intravenous glucose tolerance test (IVGTT). After model validation, we investigate how removing glucose feedback from insulin secretion, as seen in insulin-dependent diabetes, disrupts the regulation of glucose and glucagon. To do this, we simulate the model (a) when insulin secretion is reduced to mimic an insufficient dose of insulin, (b) when the peak of insulin action is delayed mimicking a dosing delay of insulin, and (c) when both occur simultaneously. Lastly, we test different half-lives of insulin to evaluate how an increased half-life of manufactured insulin may further disrupt the system. We find that when insulin secretion is decreased, glucagon still responds to high glucose levels by decreasing glucagon production. This suggests that in cases of type 2 diabetes, where glucagon secretion is elevated despite high levels of glucose, a lack of insulin response may not be the sole cause for glucagon dysfunction. We also find that delaying insulin secretion increases the risk of a hypoglycemic event through a suppression of glucagon production. Initially, the spike in glucose causes glucagon secretion to be reduced; this is then followed by the delay in insulin peak which then continues to suppress glucagon despite blood glucose levels falling, leading to a lack of response by glucagon and a subsequent hypoglycemic event. Furthermore, we find that a higher half-life of insulin causes it to remain longer in the blood stream, inhibiting glucagon’s response to severely low glucose levels (glucose levels less than 3.9 mmol/L). This sheds light on why patients taking exogenous insulin, which has a longer half-life than endogenous insulin, may have difficulty recovering from hypoglycemic events. Hence, our model suggests that keeping the half-life of exogenous insulin below 10 minutes and administering it immediately after meals could help reduce the risk of hypoglycemic events in patients with type 1 or insulin dependent diabetes. Overall, we highlight how a disruption in the feedback between insulin and glucose not only alters blood glucose levels, but also glucagon response, which may lead to further disruption of the system.

## 1. Introduction

Blood glucose levels are a crucial component of homeostatic regulation. Therefore, maintaining healthy blood glucose levels is critical to health. The regulation of glucose levels in the body is accomplished through a complex interplay of hormones, primarily insulin and glucagon, secreted by the pancreas. These hormones work together to ensure that blood glucose levels remain within a narrow and well-defined range. The significance of maintaining healthy glucose levels cannot be overstated, as levels outside a healthy range can have severe consequences. Elevated blood glucose levels, known as hyperglycemia, can lead to a range of complications such as cardiovascular diseases, kidney dysfunction, and nerve damage [22, 32]. In extreme cases, uncontrolled hyperglycemia can lead to diabetic ketoacidosis (DKA) which is life-threatening and is the leading cause of death in children with type 1 diabetes [47]. Similarly, low blood glucose levels, or hypoglycemia, also pose significant risks. Glucose is the primary energy source for the brain and other vital organs. When blood glucose levels drop too low, it can result in symptoms such as dizziness, confusion, and, in severe cases, loss of consciousness. In situations where the body cannot access an adequate supply of glucose, the consequences can be fatal [2].

The pancreatic islets of Langerhans are cell clusters found in the pancreas that respond to glucose levels by secreting either a blood glucose lowering hormone, insulin, or a blood glucose raising hormone, glucagon [34]. Alpha (*α*) cells, beta (*β*) cells, and delta (*δ*) cells are the primary cells composing the pancreatic islets of Langerhans (Figure 1), each of which respond to both increases and decreases in blood sugar levels differently [34]. For simplicity, we do not include *δ* cell dynamics in our model since glucagon and insulin are sufficient to capture data utilized in this work (see Section 4.1).

**Figure 1:**
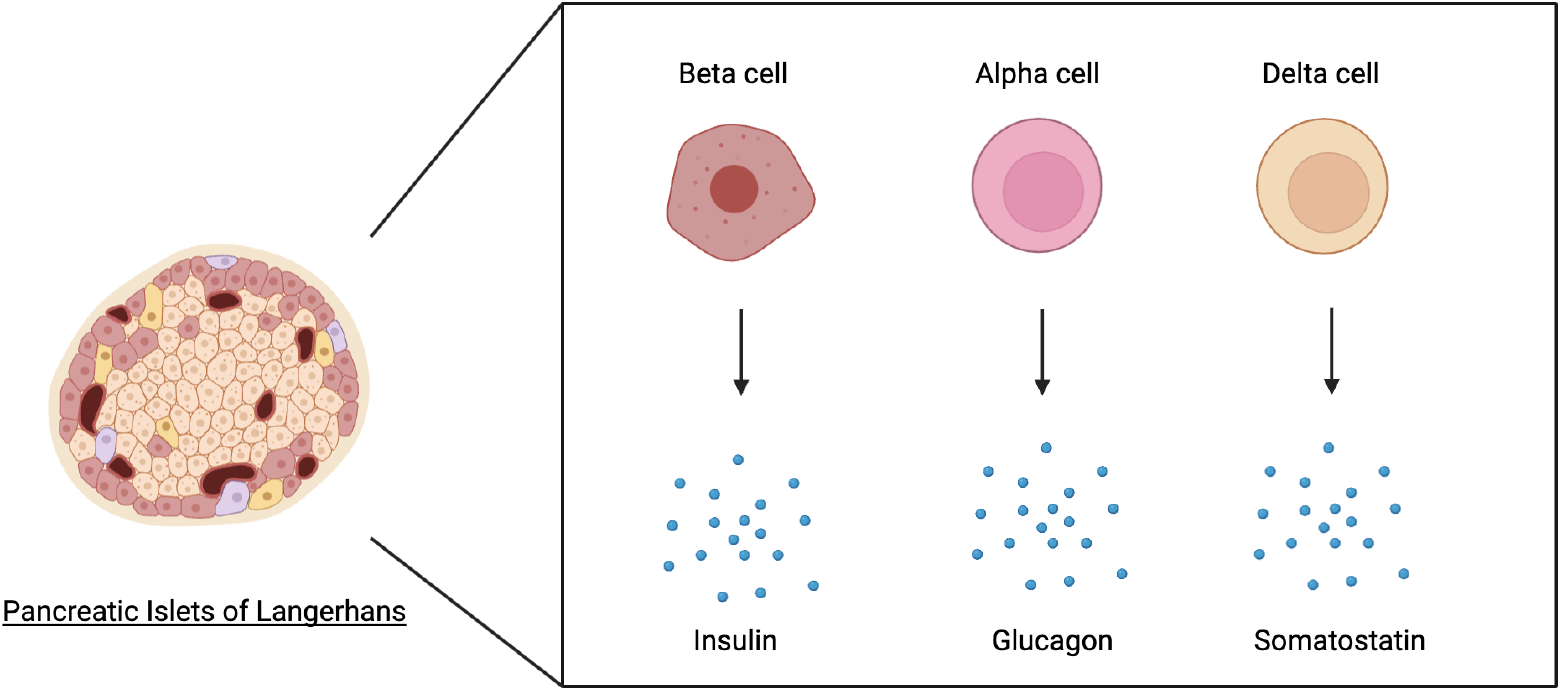
Pancreatic islet of Langerhans and the hormones secreted by *α* cells, *β* cells, and *δ* cells. [45].

To raise blood glucose levels, the pancreatic *α* cells secrete the hormone glucagon, also known as anti-insulin, to raise blood glucose levels. Glucagon works by triggering the liver to convert stored glucose (glycogen) into glucose, which is then released into the bloodstream; this process is known as glycogenolysis [23]. Moreover, glucagon can also raise blood glucose levels by preventing the liver from taking in and storing glucose so that more glucose remains in the blood stream [23]. To lower blood glucose levels, the pancreatic *β* cells secrete the hormone insulin to lower blood glucose levels. Insulin works by acting as a “key” that allows glucose to enter into a cell and subsequently lowers blood glucose levels. Without insulin, glucose cannot enter the cell [10]. Moreover, insulin blocks the production of glucagon from *α* cells, which leads to less production of glucose by the body [3].

In 1979 Richard Bergman and colleagues introduced a model of the insulin-glucose regulatory system [6] to help understand data from an intravenous glucose tolerance test (IVGTT). The test involves an injection of glucose into the blood stream followed by monitoring of blood glucose and insulin levels for a period of time. The test can help in the diagnosis of type 2 diabetes by calculating an individual’s insulin sensitivity - a measurement of the ability of insulin to stimulate the uptake of glucose.

This model was coined the *minimal model*, as it was the simplest quantitative biological framework that was able to successfully capture insulin-glucose longitudinal data [12]. However, while this model is still utilized as of writing, it does possess limitations. Two of the main criticisms of the minimal model are that (a) insulin production is dependent on time, making the system non-autonomous and (b) a non-observable remote insulin compartment is introduced that aims to model the insulin-dependent uptake of glucose [33]. Beyond the minimal model, there have been a variety of other models that aim to describe insulin-glucose dynamics [41, 26, 33, 11]. We note that a number of these models aim to remove the time dependence and remote insulin compartment from the model; however, this often leads to highly complex and overparameterized (i.e. nonidentifiable) systems and the appeal of the minimal model being minimal while still capturing the dynamics of the system should not be be overlooked. Moreover, in alignment with the growing biological research on glucagon, additional models have been introduced to explore the relationship between insulin, glucose, and glucagon [41, 16, 39, 48]. However, many of these models are derived from the minimal model, which results in the aforementioned criticisms.

Efforts to understand the role of *α* cells in secreting glucagon have recently started to gain traction, as the role of glucagon in maintaining glucose homeostasis is more well understood [27, 17]. Some sources suggest that *α* cells are primarily regulated by neighboring *β* cells, while others contend that *α* cells possess intrinsic mechanisms for detecting and responding to glucose [25, 8]. Moreover, research has suggested that glucagon may play a role in the development of both type 1 and type 2 diabetes [28, 21]. More generally, there is also research investigating the role of glucagon in regulating blood glucose levels, as well as possible mechanisms that may cause glucagon dysfunction in both type 1 and type 2 diabetes [25]. In this work, we develop a model of the insulin-glucose regulatory system that is self-regulatory and includes a direct interaction between insulin and glucose (i.e.there is no remote insulin compartment). Furthermore, we add glucagon as a third compartment of our model. In fact, we note that the addition of glucagon is necessary to fit to time course data of insulin and glucose, highlighting an important role for glucagon in insulin-glucose dynamics. The model is presented in Section 2, and results of model fitting can be seen in Section 4.1; Section 3 contains methods related to the (publicly available) experimental data incorporated into the mathematical model, as well as parameter estimation techniques. After model validation, we investigate in Section 4.3 how removing glucose feedback from insulin secretion, as is seen in insulin-dependent diabetes where exogenous insulin is administered, disrupts the regulation of glucose and glucagon. To do this, we simulate the model (a) when insulin secretion is reduced to mimic an insufficient dose of insulin, (b) when the peak of insulin action is delayed mimicking a dosing delay of insulin, and (c) when both occur simultaneously. Lastly, we varied the half-life of insulin to evaluate how an increased half-life of manufactured insulin (when compared with endogenous insulin) may further disrupt the system. We found that when insulin secretion is decreased, glucagon still responds to high glucose levels by decreasing glucagon production (Figure 8a). Next, we found that delaying insulin secretion increases the risk of a hypoglycemic event through a suppression of glucagon production (Figure 8b). Lastly, we found that the higher the half-life of insulin, the longer it remains in the blood stream, inhibiting glucagon’s response to severely low glucose levels (glucose levels less than 3.9 mmol/L; Figure 9). Overall, we highlight how a disruption in the feedback between insulin and glucose does not only alter blood glucose levels but also glucagon response, leading to further disruption of the system.

## 2. Model

We develop a three compartment model of insulin and glucose dynamics that includes insulin, glucose, and glucagon. Mass action kinetics are assumed for most interactions, unless otherwise specified. The major biological features of the model are the following:

1. Glucose stimulates the production of insulin [14]. The production of insulin was assumed to saturate as a function of glucose [12]. That is, the maximum insulin production rate (*k*_1_) is independent of glucose.
2. Insulin regulates the insulin dependent uptake of glucose (*δ*_2_ *I*) which is the primary mechanism by which glucose enters cells [13].
3. A small amount of glucose can enter cells through an insulin independent uptake of glucose (*δ*_3_) [13].
4. The production of glucagon is inhibited by both high levels of insulin (*p I*) and glucose (*z*(*G*− *h*)^*+*^)[38]. The inhibition by glucose is modeled as *z*(*G* − *h*)^*+*^ due to the assumption that when G is below a baseline value *h*, glucose does not inhibit glucagon at all, as observed in the response of glucagon to low glucose levels. Thus, when *G < h* glucose no longer inhibits glucagon production. For *x* ∈ ℝ, recall that *x*^*+*^ denotes the positive part of *x*:

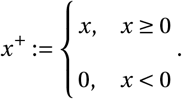
5. The production of glucose in the body (that is, not due to a glucose injection) is regulated by glucagon (*k*_3_*G*_*𝓁*_) [18]. This occurs via glucagon signaling to the liver to break down stored glucose and releasing it into the blood stream [23].
6. Insulin and glucagon both decay naturally (*δ*_1_ and *δ*_4_, respectively) [37].

Under these assumptions, the model schematic of the three compartment insulin-glucose-glucagon model is shown in Figure 2. A system of three ordinary differential equations incorporating the above assumptions is given below:

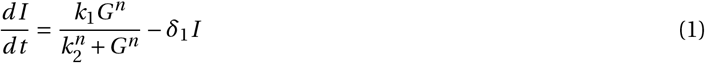

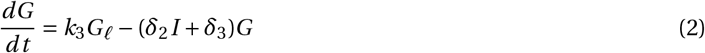

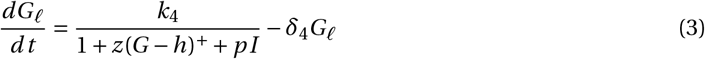

**Figure 2:**
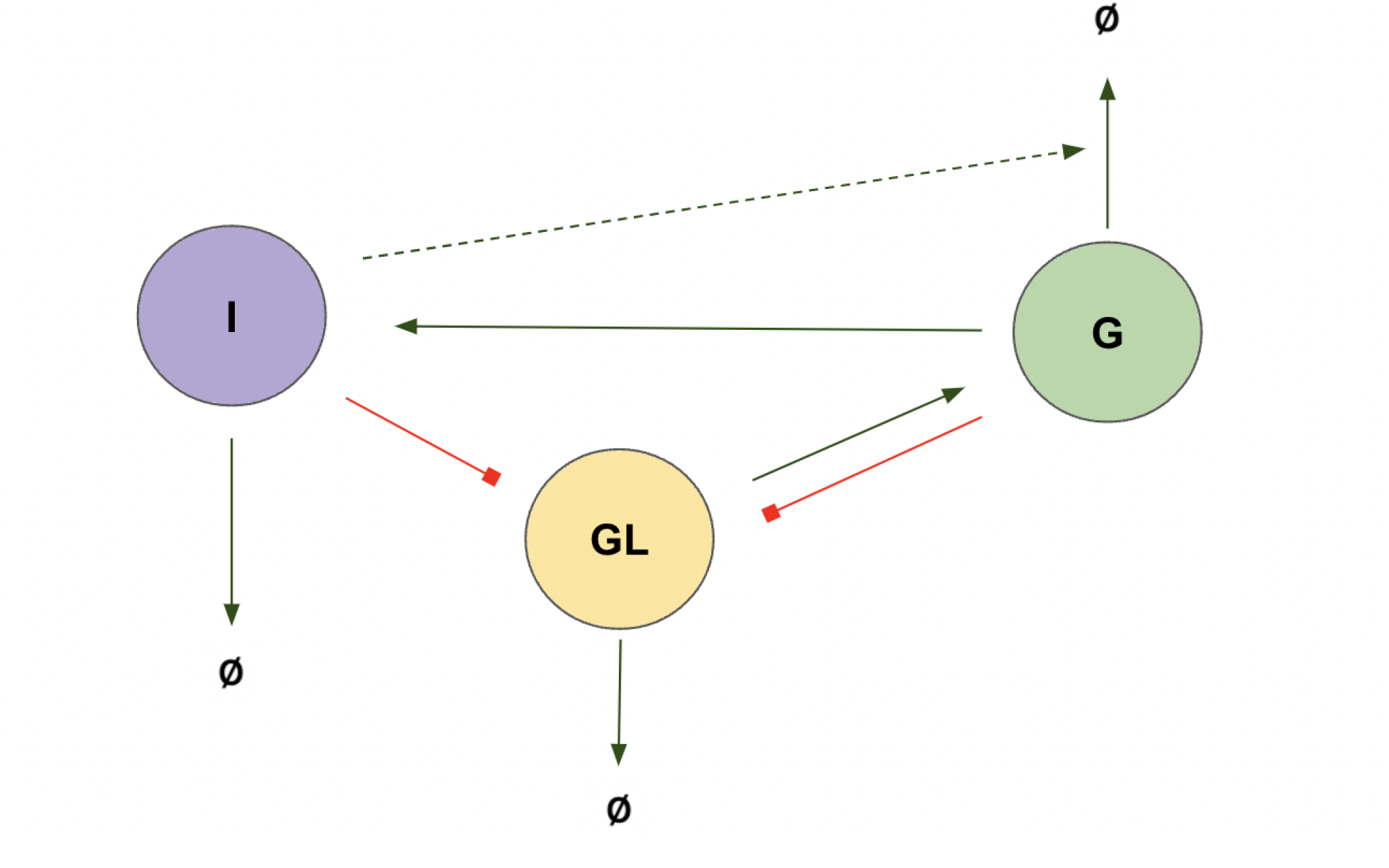
Three compartment model schematic representing insulin-glucose-glucagon model (1)-(3).

It can be shown that the above system possess a unique positive steady state (see Section 7.1 in the Supplementary Material). We note that we have also developed a two compartment model of insulin and glucose alone to assess the necessity of glucagon in explaining insulin-glucose dynamics; see Section 7.2 in the Supplemental Material. Results suggest that three compartments are necessary to fully capture the utilized experimental data (see Figure 11).

To simulate a glucose injection from the IVGTT, we incorporate an external input *u*(*t*) to equation (2) of the form

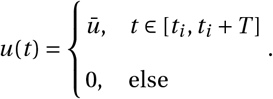

Here, 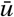 is the applied (constant) dosage which was taken to match the peak glucose value of the experimental data (see Table 1 for value), *t*_*i*_ is start time of the dosage, and *T* is the length of infusion. In this case, system (1)-(3) takes the form

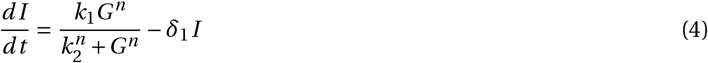

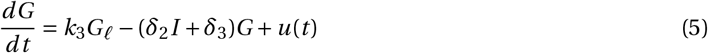

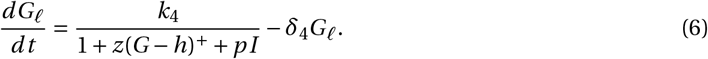

**Table 1:** Table of initial estimate ranges, best fit parameter values, and units for the insulin-glucose-glucagon model 1 - 3.

| Parameter | Initial Range<br>(multi-start) | Estimated<br>Parameter Value | Units |
| --- | --- | --- | --- |
| $k_1$ | [0 2] | 0.13 | $\frac{\mu g}{mL \cdot mins}$ |
| $k_2$ | [0 100] | 9.81 | $\frac{mmol}{L}$ |
| $k_3$ | [0 2] | 0.01 | $\frac{mmol}{pmol \cdot mins}$ |
| $k_4$ | [0 2] | 21.9 | $\frac{pmol}{L \cdot mins}$ |
| $\delta_1$ | [0 2] | 0.13 | $\frac{1}{mins}$ |
| $\delta_2$ | [0 2] | 0.17 | $\frac{mL}{\mu g \cdot mins}$ |
| $\delta_3$ | [0 2] | 0.01 | $\frac{1}{mins}$ |
| $\delta_4$ | [0 2] | 0.35 | $\frac{1}{mins}$ |
| $p$ | [0 10] | 99.87 | $\frac{mL}{\mu g}$ |
| $z$ | [0 10] | 1.35 | $\frac{L}{mmol}$ |
| $h$ | [0 4] | 2.45 | $\frac{mmol}{L}$ |
| $n$ | [0 10] | 5.64 | - |

**Table 2:** Table of initial estimate ranges, best fit parameter values, and units for the insulin-glucose model (32) - (33).

| Parameter | Initial Range | Best Fit Parameter Value | Units |
| --- | --- | --- | --- |
| $k_1$ | [0, 1] | 0.22 | $\frac{\mu g}{mL \cdot minutes}$ |
| $k_2$ | [0, 1] | 11640 | $\frac{mmol}{L}$ |
| $k_3$ | [0, 1] | 0.04 | $\frac{mmol}{L \cdot minutes}$ |
| $\delta_1$ | [0, 1] | 0.11 | $\frac{1}{minutes}$ |
| $\delta_2$ | [0, 10] | 0.28 | $\frac{mL}{\mu g \cdot minutes}$ |
| $\delta_3$ | [0, 2] | 0.0001 | $\frac{1}{minutes}$ |
| $n$ | [0, 1] | 3.36 | - |

## 3. Methods

### 3.1. Experimental Data

We utilized data collected from a study conducted by Manell et al. [29], where ten pigs were subjected to an Intravenous Glucose Tolerance Tests (IVGTT). These pigs underwent measurements of glucose, insulin, and glucagon, which were taken both before glucose infusion and at 2, 5, 10, 20, 30, 45, 60, 90, 120, and 180 minutes post infusion. Of the ten pigs, the final glucagon measurement was missing for three of them.

We fit the model (4)-(6) to the median (with respect to the ten pigs) temporal data. We used the median to avoid outliers skewing the estimated parameter values, particularly with respect to glucagon, where the measurements exhibited a large variance (see Figure 3). Furthermore, we computed a 95% confidence interval with respect to the median via bootstrapping [1]. Specifically, we generated 10,000 new data sets, with each set consisting of 10 pigs. We then calculated the median of each new data set to generate 10,000 medians which gave a distribution of the medians at each time point. From this data we constructed a sampling distribution for the median, which was used to calculate the 95% confidence interval. The longitudinal pig data, together with the median and estimated confidence intervals are provided in Figure 3. We highlight that while glucose and insulin exhibited minimal variation over time, the levels of glucagon displayed substantial variability, particularly during the 30 minutes following the IVGTT.

**Figure 3:**
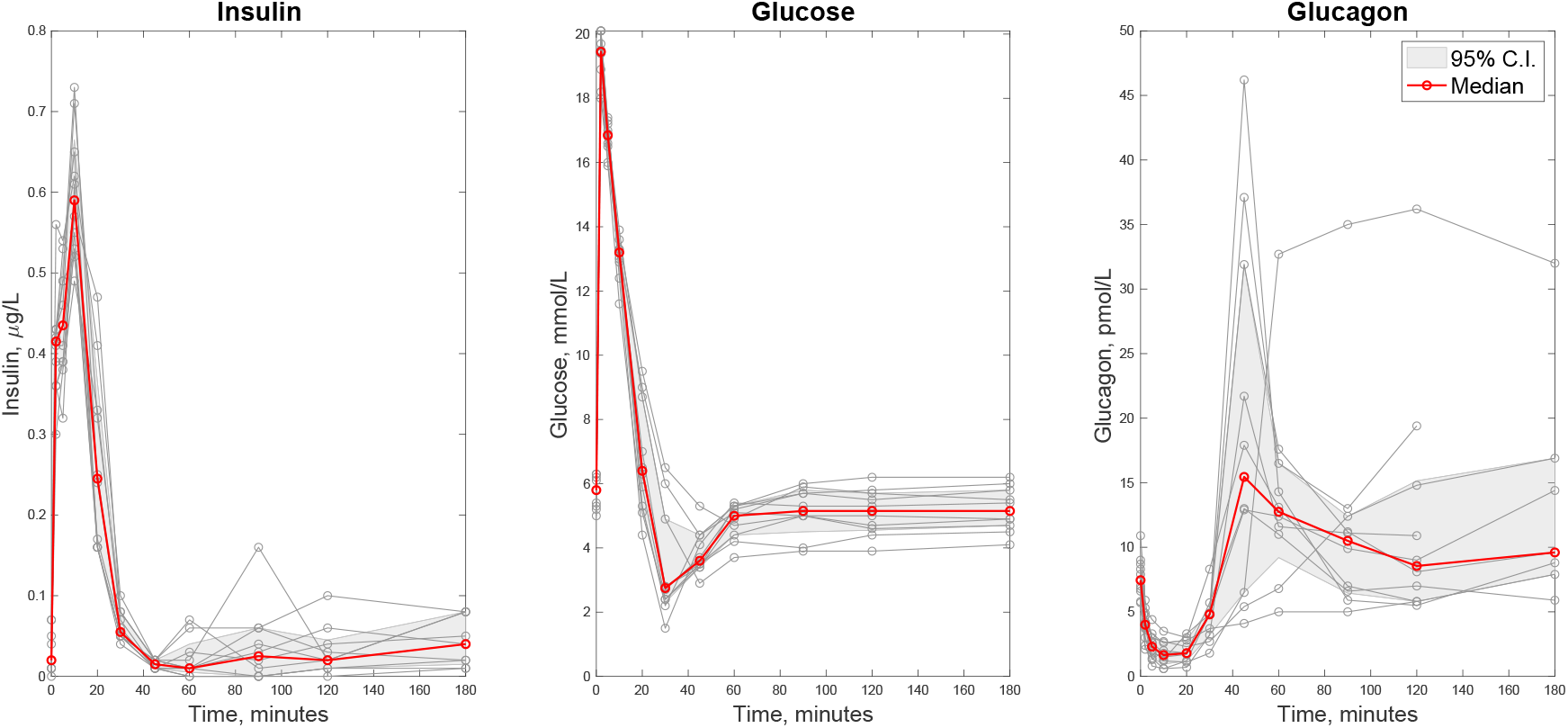
Experimental Data. Plot of the pig data (insulin, glucose, and glucagon over a 180 minute period), medians, and estimated 95% confidence intervals for an IVGTT. Left is insulin, middle is glucose and right is glucagon.

### 3.2. Parameter Estimation

To estimate rate parameters in 1-3, we fit to the median of the pig data presented in Section 3.1. The median of the pig data is measured at times *t*_*i*_, *i =* 1, 2,…, *n*; thus the experimental data consists of tuples of the form 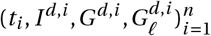. For example, *G*^*d, i*^ denotes the median glucose level at time *t*_*i*_.

We utilize a normalized least-squares inspired cost functional which penalizes deviations from all three compartments (insulin, glucose, and glucagon). Specifically, we define

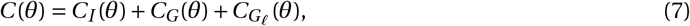

where

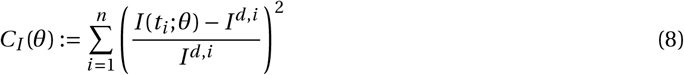

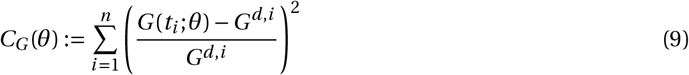

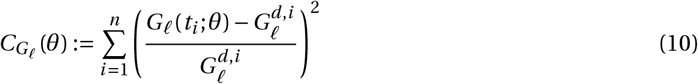

Here *I* (*t*_*i*_; *θ*) denotes the solution of the insulin compartment of the ODE system 1-3 evaluated at time *t*_*i*_, as a function of the model parameters *θ*; similar statements holds for *G*(*t*_*i*_, *θ*) and *G*_*𝓁*_(*t*_*i*_, *θ*). The parameter vector

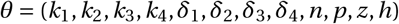

is then estimated to minimize *C*, with respect to non-negativity constraints, utilizing *fmincon* in Mat-lab’s nonlinear optimization toolbox. We note that when calibrating the minimal model to the same data set, glucagon is not explicitly considered (see equations 12-14), and thus the following analogous cost functional is utilized:

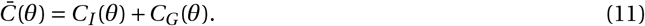

Multi-start optimization was performed, and the initial parameter ranges utilized are provided in Table 1. We note that while the initial conditions for the optimization algorithm are contained in this range, the best fit parameter value may lie outside of it. To ensure that a global minimum was achieved, we repeated the optimization 1000 times while randomizing initial estimates between the ranges seen in Table 1. We used an *n*-dimensional Sobol set transformed to the hypercube [0, *γ*]^*ρ*^ where *ρ* is the length of the parameter vector *θ* and *γ* is the upper bound for each parameter value. During optimization, all parameter values were constrained to remain non-negative.

After the multi-start optimization was performed, we constructed tornado plots to ensure that we located the global minimum of the cost function. The tornado plots were generated by varying one parameter at a time within 10% of its computed optimal value, and subsequently calculating the cost functional value. If a change in parameter value led to a decrease in the cost functional, the parameter value was changed accordingly, and the process was repeated until all local changes in parameter values led to only increases in the cost 5.

## 4. Results and Discussion

### 4.1. Model fitting

We begin by estimating parameters in the insulin-glucose-glucagon model ((1)-(3)) by fitting the model to the data presented in Section 3.1, as discussed in Section 3.2; the code to generate all figures can be found on github. The optimal parameter set is defined to be the one that minimizes the cost functional defined in equation (7). The results are provided in Figure 4. We see that our model is able to capture the median of the data well. The cost functional at the optimal parameters is *C =* 0.96976, with an adjusted *R*^2^ value of 0.9283.

**Figure 4:**
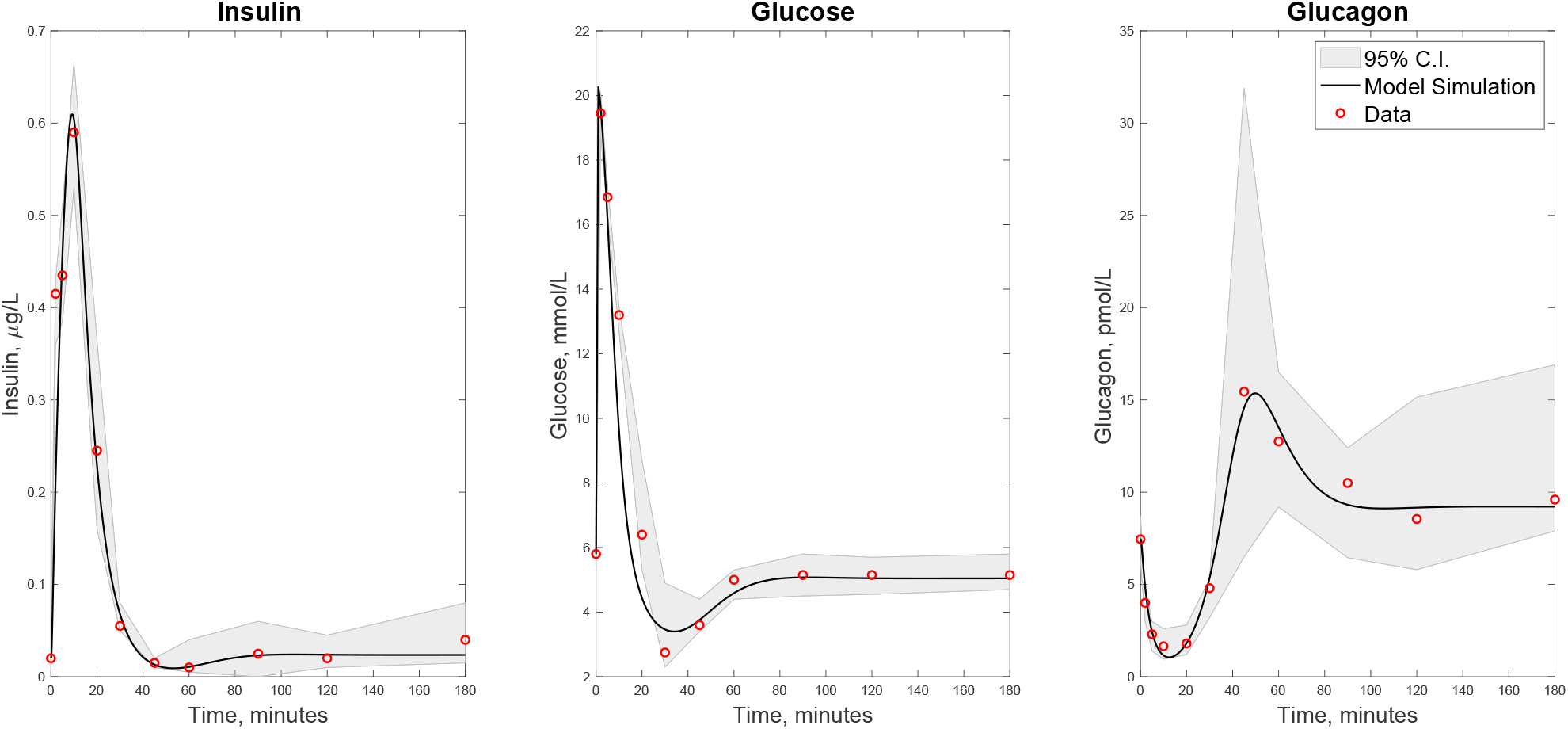
Model Fits. The best fit for the insulin-glucose-glucagon model 1-3. Here, the constant dosage 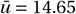 was used with initial time *t*_*i*_ *=* 0 and dose length *T =* 1. The initial conditions for insulin, glucose and glucagon were taken to be the initial values of the data. The cost functions evaluated at the optimal set of parameters are *C =* 0.96976 (insulin-glucose-glucagon cost) and 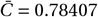 (insulin-glucose cost). See Table 1 for the corresponding estimated parameters. Left is insulin, middle is glucose and right is glucagon.

The estimated parameter values are provided in Table 1. Figure 5 also shows tornado plots when individually varying each best fit parameter within 5%, 10%, 15%, and 20% of its original value. Note that varying each parameter does not lead to a decrease in the cost function *C*, providing evidence that we are indeed at a global minimum.

**Figure 5:**
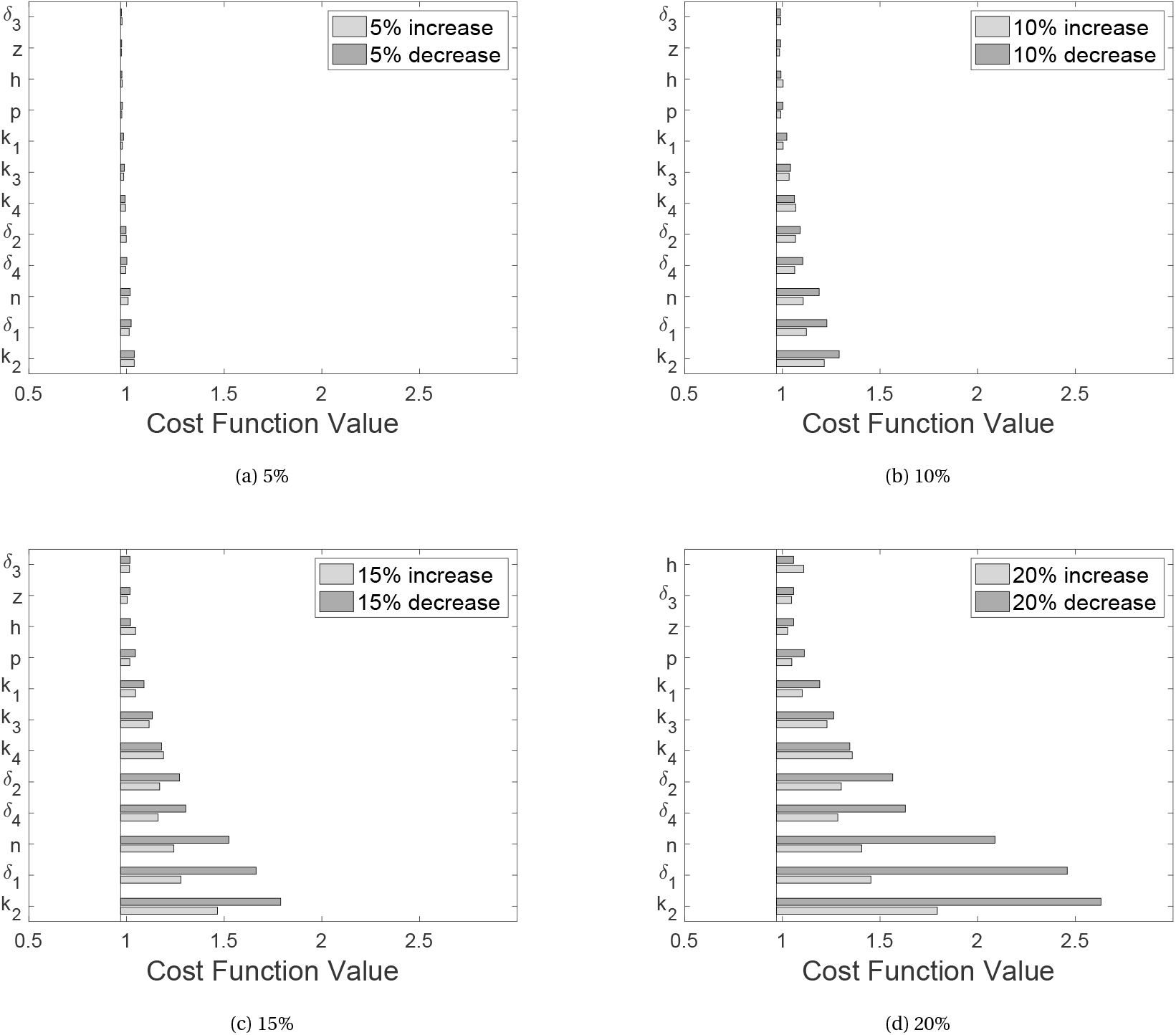
Local Minimum Validation. Tornado plots for the change in the cost function (*C*) when varying each parameter by the given percentage of its optimal value. The vertical line represents the minimum residual. Parameter values in (a) are changed by 5%, (b) are changed 10%, (c) are changed by 15% and (d) are changed by 20%.

For comparison, we also fit the minimal model developed by Bergman [4] to the same data set. The minimal model consists of three compartments: insulin (*I*), glucose (*G*) and remote insulin (*X*). Here, insulin does not directly inhibit glucose; rather insulin enters a remote compartment (*X*) and *X* acts to inhibit glucose. Insulin production is a function of both time and the degree to which glucose exceeds a baseline threshold value *h*. Glucose production is constant with respect to a baseline level of glucose, which is dependent on the specific data set. More precisely, *G*_*b*_ is taken to be the initial value of glucose prior to the start of the IVGTT. Note that *G*_*b*_ is not estimated from the data. The production of remote insulin depends on *I*, and glucose uptake occurs through both an insulin dependent and independent pathway. Lastly, both insulin and remote insulin have natural decay rates *n* and *P*_2_, respectively. The differential equations describing these dynamics are provided below; for more details on the model formulation, see [4].

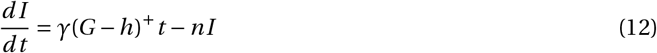

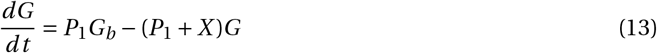

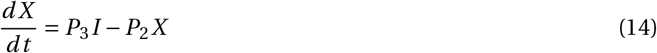

Figure 6 shows the result of the best fit of the minimal model (12)-(14) to the same data set utilizing the procedure outlined in Section 3.2 compared with the best fit for the insulin-glucose-glucagon model. Recall that when estimating parameters in the minimal model, we consider the cost function (11). The cost 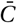 for the minimal model is 1.5 whereas cost 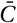 for the insulin-glucose-glucagon model is 0.78. To adjust for the insulin-glucose-glucagon model possessing a larger number of parameters, we also calculated the adjusted *R*^2^ value for each model; the adjusted *R*^2^ for the minimal model is 0.77, while that of the insulin-glucose-glucagon model is 0.92. The best fit parameter values and tornado plots for the minimal model can be found in the Supplemental Material (see Section 7.3, Table 3, Figure 14).

**Table 3:** Table of initial estimate ranges, best fit parameter values, and units for the minimal model (12) - (14).

| Parameter | Initial Range | Best Fit Parameter Value | Units |
| --- | --- | --- | --- |
| $\gamma$ | [0, 1] | 0.77 | $\frac{L\mu g}{mmolmLtime^2}$ |
| $h$ | [0, 1] | 3.98 | $\frac{mmol}{L}$ |
| $n$ | [0, 1] | 100.35 | $\frac{1}{time}$ |
| $P_1$ | [0, 1] | 0.01 | $\frac{1}{time}$ |
| $P_2$ | [0, 10] | 0.59 | $\frac{1}{time}$ |
| $P_3$ | [0, 2] | 0.24 | $\frac{mmol}{L}$ |
| $G_b$ | - | - | $\frac{mmol}{L}$ |

**Figure 6:**
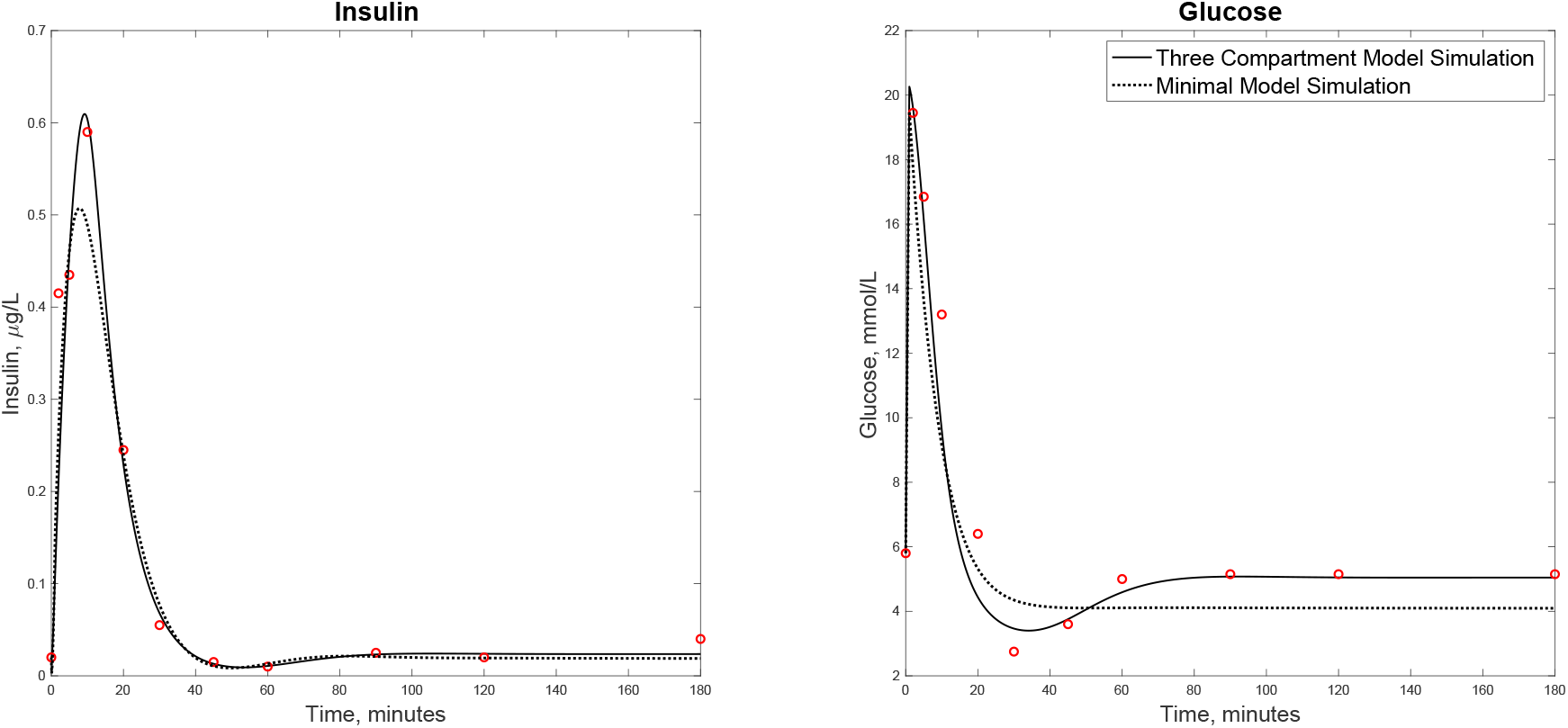
Minimal Model Fits. The best fit for the minimal model (12) -(14) together with the best fit of the three compartment model (1) - (3). The cost function at the given parameters for the minimal model fit is 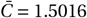, with a corresponding *R*^2^ value of 0.7569. Left is insulin and right is glucose.

### 4.2. Insulin Sensitivity and Glucose Effectiveness

One of the main applications of the minimal model is its ability to provide a patient’s glucose effectiveness and insulin sensitivity to both aid in diagnosing a patient with type 2 diabetes, as well as understanding the severity of the disease [43, 15].We now explore how to calculate both insulin sensitivity and glucose effectiveness using our three compartment insulin, glucose and glucagon model (1) - (3). We utilize Bergman’s definitions to calculate both glucose effectiveness and insulin sensitivity. Specifically, Bergman defines glucose effectiveness (*E*) as the quantitative enhancement of glucose disappearance due to an increase in the plasma glucose concentrations [5]. In other words, glucose effectiveness is the ability of glucose to stimulate its own uptake and is defined quantitatively as follows:

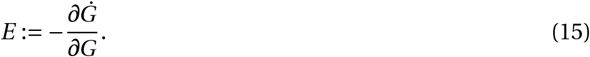

Here *Ġ* is the rate of change of the plasma glucose concentration *G*, i.e. effectiveness for model (1) - (3) is given by *Ġ = dG*/*dt*. Thus, glucose effectiveness for model (1) - (3) is given by

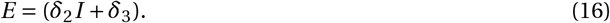

Bergman also defines the insulin sensitivity (*S*) as the quantitative influence of insulin to increase the enhancement of glucose disappearance [5]. That is, insulin sensitivity is the ability of insulin to stimulate glucose disappearance and is given by the following formula:

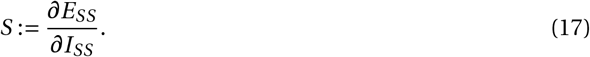

Here the subscript *SS* indicates that the quantities are measured at steady state. Using the above definition, insulin sensitivity for model (1) - (3) is

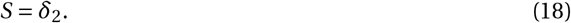

The insulin sensitivity factor is easy to interpret as *δ*_2_ is the parameter that controls the impact of insulin on glucose clearance. A high *δ*_2_ value would result in an increased clearance of glucose and indicate sensitivity to insulin, whereas a low *δ*_2_ would result in less clearance of glucose and indicate insulin resistance [24]. Our insulin sensitivity factor estimated by fitting the median of the pig data is 0.17. In [5], Bergman estimates insulin sensitivity factors between 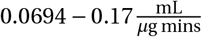 (using the conversion factor 1*µU =* 0.0347*µg*; [20]). Hence, our insulin sensitivity factor appears to be biologically reasonable.

### 4.3. Removing Glucose Feedback on Insulin

Diabetes is a disease where the body either becomes resistant to insulin (type 2 diabetes) or there is a low/absent production of insulin due to an autoimmune attack on the insulin producing *β* cells (type 1 diabetes). In the latter case, and occasionally in the former as well, the patient requires insulin injections in order to regulate blood glucose levels which results in the loss of direct feedback of glucose on insulin production. This loss of communication has a large impact on the insulin-glucose regulatory system; indeed, the impact is significant to the degree so that glucagon no longer functions properly despite *α* cells remaining functional [7]. Understanding the disruption in glucagon dynamics caused by insulin disruption may be critical to better understanding the role of glucagon both before and after the diagnosis of diabetes. In this section, we investigate this loss of glucose feedback on both glucose and glucagon dynamics using a simplified system derived from (1) - (3).

A schematic of the considered system with no glucose feedback on insulin is provided in Figure 7. We are thus analyzing a system where the communication between insulin and glucose is disrupted, so that insulin can be considered an external input, which operates independently of glucose (*G*) and glucagon (*G*_*𝓁*_). For clarity, we denote this external insulin concentration with the symbol *Î = Î* (*t*). This leads to the following two-dimensional system with external input *Î*:

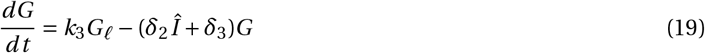

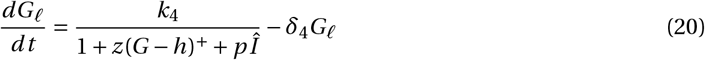

**Figure 7:**
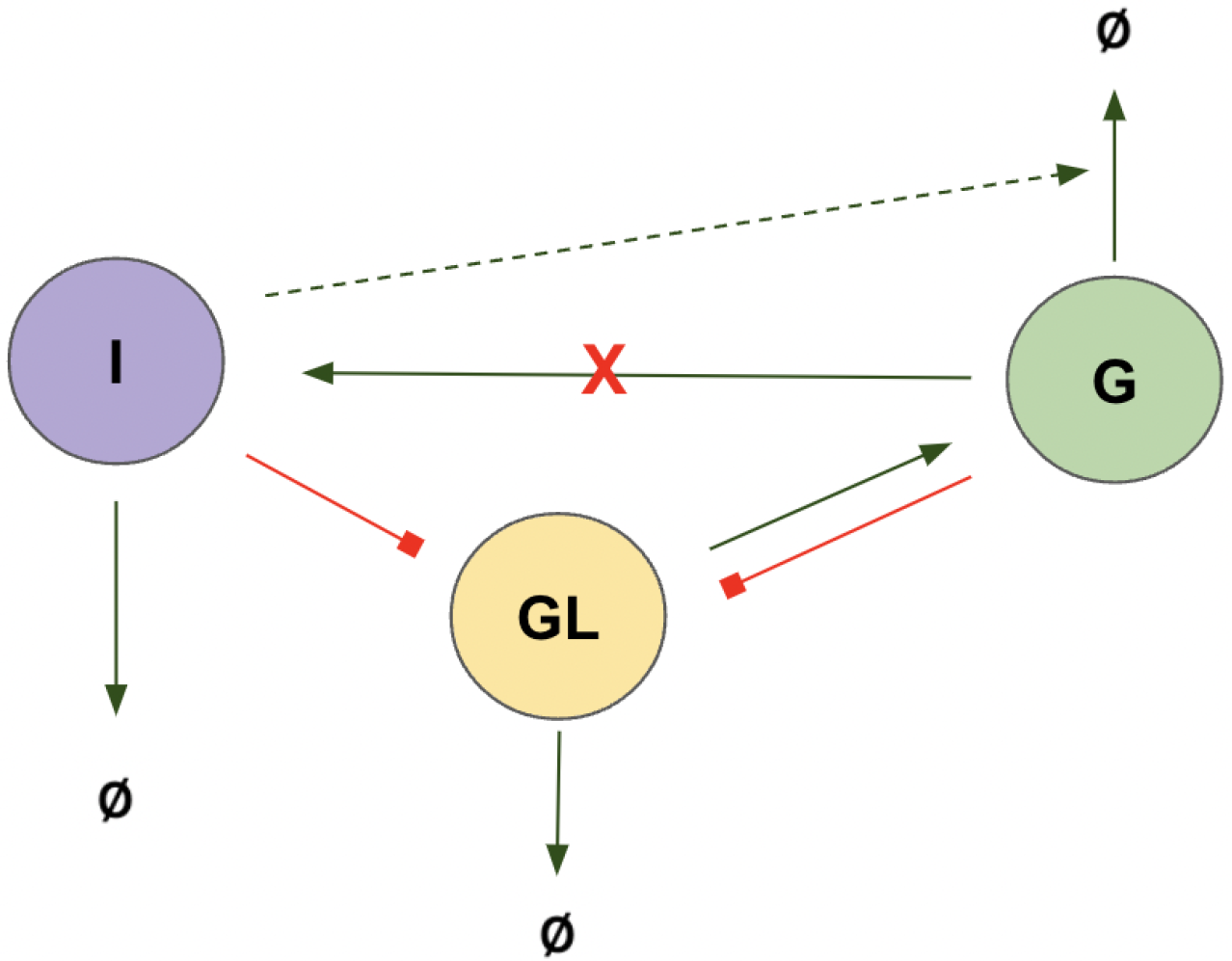
Schematic of the insulin-glucose-glucagon system with glucose feedback on insulin removed. The red *X* denotes that the interaction is not considered in system (19) - (20), and insulin is considered as an external input to the system (*Î*).

**Figure 8:**
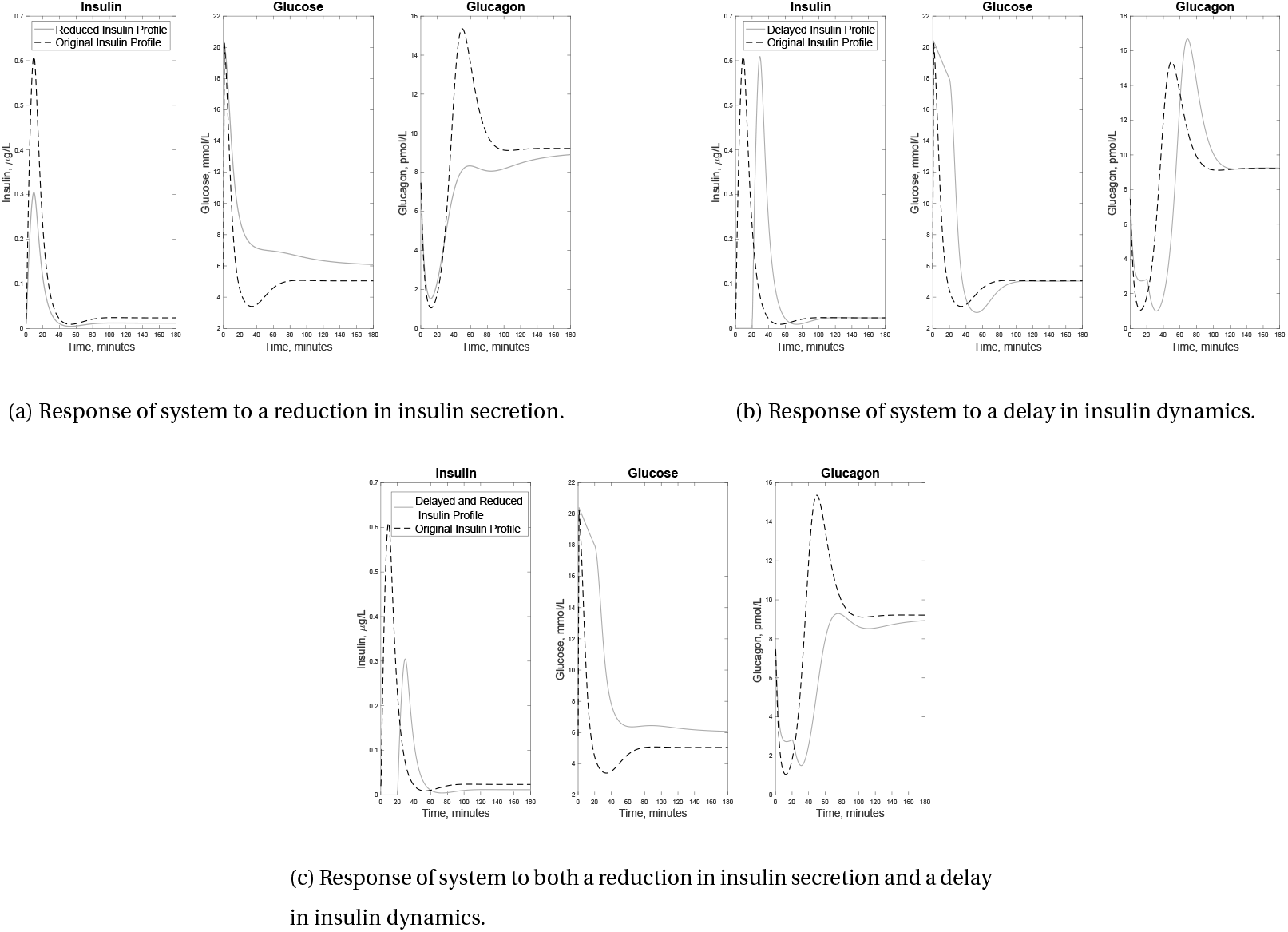
Dynamics in the Absence of Glucose Feedback. Dynamics of insulin, glucose, and glucagon (19) - (20) when altering the insulin profile from the model fit (Figure 4). Recall that in this system, there is no feedback of glucose on insulin. The insulin profile in (a) is reduced, in (b) is delayed and in (c) is both reduced and delayed.

**Figure 9:**
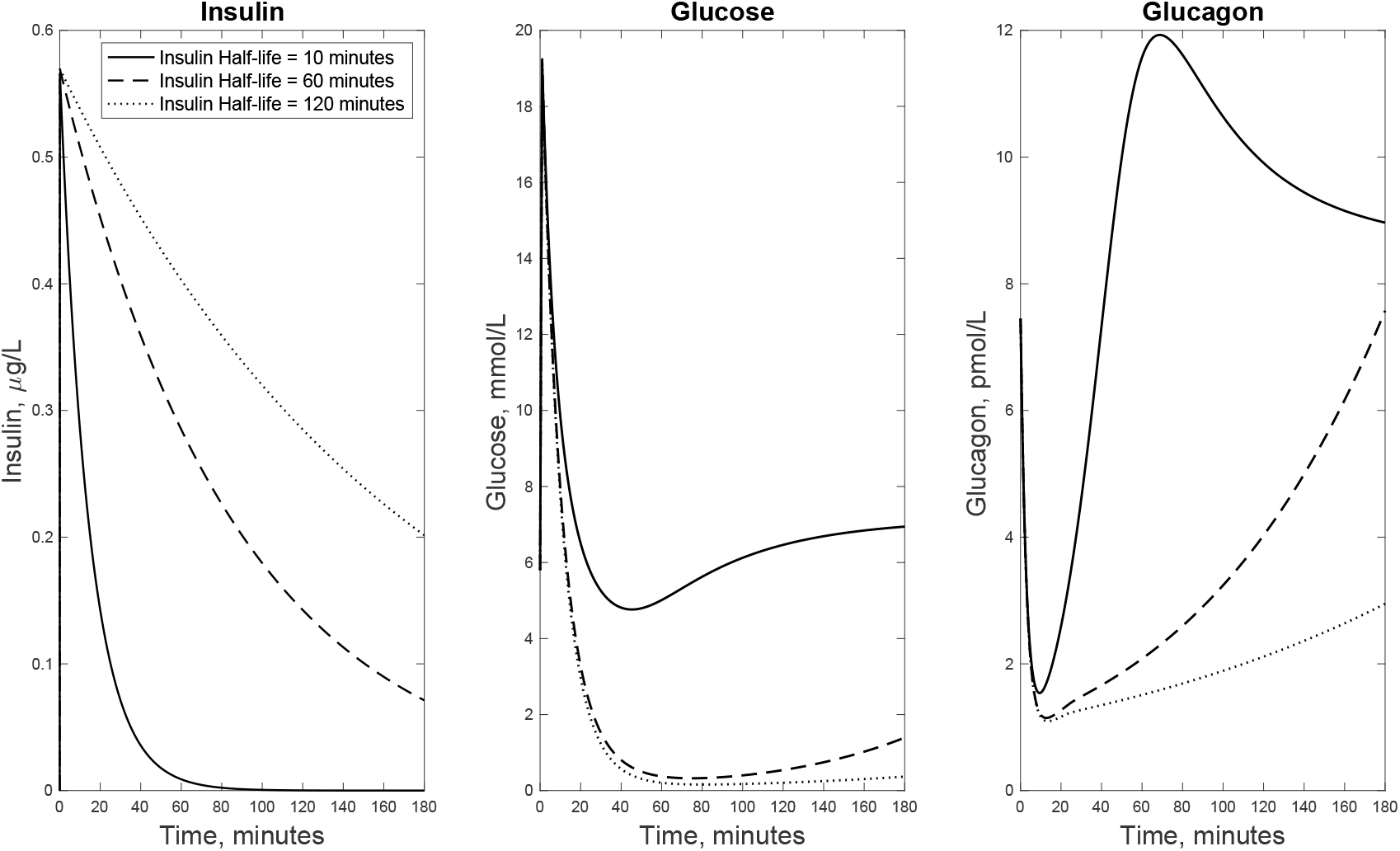
Varying Insulin Half-lives. Insulin, glucose, and glucagon dynamics in system (19) - (20) when varying the half-life of insulin to 10, 60, and 120 minutes. Insulin dynamics are given by (21). Left is insulin, middle is glucose and right is glucagon.

To investigate how the removal of glucose feedback on insulin production impacts glucose and glucagon dynamics, we utilize the fit insulin profile from Figure 4 as a baseline, and vary it in different ways to simulate simplified scenarios that occur in diabetes (specifically in type 1 diabetes). We note that in the absence of any perturbation to the insulin profile *Î*, the solution to system (19) - (20) produces the dynamics observed in Figure 4 precisely (i.e. to the solution of the original model system (19) - (20)).

To simulate different scenarios occurring in diabetes, we alter the insulin input in two different ways and then combine these two alterations and investigate the impact of these alterations on glucose and glucagon dynamics. First, we simulate the dynamics of glucose and glucagon when decreasing insulin secretion; specifically, we take the insulin profile from Figure 4 and reduce the total (integral) insulin concentration by 50% (Figure 8a, top left), which provides a new insulin profile *Î*. A reduction in insulin secretion can be seen in both type 1 and type 2 diabetes. In type 1 diabetes, a reduction in insulin secretion due to loss in beta cells mass may arise during disease progression. In type 2 diabetes, a reduction in insulin secretion may occur due to a decline in the function of beta cells [9].

As opposed to the case of a simple reduction in insulin, we also investigate the dynamics of glucose and glucagon when the peak insulin concentration is delayed. Here, we again take the curve associated with the insulin profile in Figure 4, and delay the insulin dynamics by 20 minutes to produce the external signal *Î* (i.e. we shift insulin dynamics by 20 minutes, Figure 8b top right). This again has applications in both type 1 and type 2 diabetes. In type 1 diabetes, patients may inject insulin after a glucose spike is already observed resulting in a delayed reaction of insulin to a glucose spike, while in type 2 diabetes, it has been shown that insulin secretion may be delayed [42]. Lastly, we combine both the reduction in insulin secretion and the time delay to create a perturbed insulin profile *Î* (Figure 8c bottom), both of which have application in type 1 and type 2 diabetes for the aforementioned reasons. The results of these three scenarios are shown in Figure 8.

Figure 8a shows the response of system (19) - (20) when the total external insulin concentration is reduced by 50%. As expected, we see that a reduction in insulin secretion leads to higher glucose levels. That is, when compared with the original insulin profile (around 20 mins), glucose levels remain higher over time; however, we note that the glucose spike is relatively constant with respect to both models (approximately 20 mmol/L), meaning that less insulin does not increase the glucose peak. On the other hand, we observe that glucagon has an increased minimum value compared to the original insulin profile, which is most likely due to the reduction in insulin peak. This is because the reduction in insulin leads to less suppression of glucagon production. Interestingly though, the maximum of glucagon is greatly reduced despite lower levels of insulin. Therefore, this reduction in glucagon production is caused by the higher blood glucose levels. This is particularly interesting because while it has been documented that glucagon secretion depends on both insulin and glucose, the role each plays is not fully understood. The results appearing in Figure 8a thus suggest that high glucose levels can greatly reduce glucagon production in the near absence of insulin. This demonstrates that the irregular patterns of glucagon secretion in type 2 diabetes (in particular, over secretion) may not be explained solely by the dysfunction of *β* cells, supporting research that has suggested the *α* cells (the glucagon-secreting cells) are also dysfunctional in type 2 diabetes [19, 31, 46].

Figure 8b shows the simulation results when delaying insulin secretion, which is relevant in type 1 diabetes in the case when insulin is administered after, as opposed to during, a meal. Here, the insulin profile *Î* is obtained by a time delay of 20 minutes compared to figure Figure 4. In this case, we now observe a significant hypoglycemic event taking place between 40 and 80 minutes post-glucose injection. Interestingly, in this simulation the maximum insulin concentration was not reduced, yet glucose levels significantly decreased when compared to the original insulin profile. This is most likely due to the dynamics of glucagon in the system. We see that glucagon initially decreases due to the rise in glucose levels. This reduction in glucagon secretion then reduces the production of glucose. Furthermore, glucagon remains suppressed due to increased insulin levels. That is, the insulin delay prevents glucagon from responding to lower glucose levels leading to a significant hypoglycemic event (*≈* 3 mmol/L). Then, when insulin begins to fall and glucagon begins to respond to lower glucose levels, we observe a larger peak in glucagon which causes glucose levels to significantly overshoot its steady state value. The responses of both glucose and glucagon to the time shift in insulin suggest that a small delay in insulin action can lead to an increased risk of hypoglycemia, an observation that is important for patients administering insulin [44]. Furthermore, these results suggest that glucagon plays an important role regulating glucose levels by reducing the production of glucose. Thus, glucagon has dual roles in the regulation of blood sugar: in both helping lower glucose levels, as well as its primary role in raising blood glucose.

Figure 8c shows simulation results when combining both a reduction and a delay of the insulin profile. Note here that glucose levels return close to baseline values without a hypoglycemic event although they remain elevated for a longer period of time. Moreover, the decrease in glucose levels is delayed when compared with the original simulation. Glucagon dynamics closely resemble the original glucagon dynamics, with the exception of the time delay and a reduction in the peak. These results suggest that when insulin is administered later, less insulin may be needed to avoid hypoglycemic events, which is most likely due to the proper regulation of glucagon.

In the feedback system (1) - (3), insulin and glucagon interact synergistically to regulate glucose levels. However, when feedback on insulin is removed, a time delay in insulin results in disruption to glucagon dynamics, and thus leads to impaired regulation of glucose. In order to optimize glucagon’s regulatory role in this case, our simulation suggest it would be best to time insulin administration so that the insulin peak and glucagon minimum occur at similar times, which seems to occur directly following a glucose spike. In fact, delays in insulin administration leading to low glucose levels is a fairly well documented phenomenon, particularly in type 1 diabetes [40, 35]

As a final application, we consider the response of the system to doses of insulin with different half-lives. The aim of this is to more closely resemble patients with diabetes, as insulins that are administered have a longer half-life compared to endogenous insulin [36]. We consider a simple exponentially decaying external insulin profile *Î*, with decay rate *k* and administered (constant) dosage *U*. The dose was assumed to be given at *t =* 0.01 minutes after the glucose injection, so that

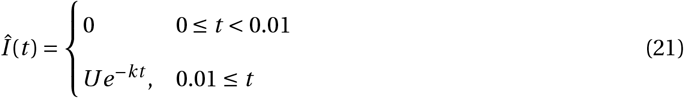

Recall that the decay rate is related to the half-life *τ* via

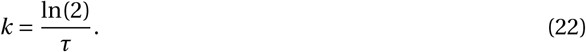

The dose *U* was taken to be the maximum of the median insulin data minus the initial value of the median insulin data to closely resemble the endogenous insulin response upon glucose injection. That is,

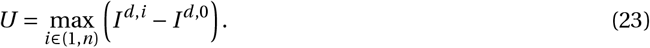

We considered three possible half-lives: 10 minutes to closely replicate endogenous insulin, 60 minutes to replicate a majority of meal time FDA approved insulins, and 120 minutes to investigate the effects of longer-lasting insulin. Note here that insulin was taken to be an intravenous dose rather than a sub-cutaneous dose for simplicity (i.e. pharmocokinetics are ignored). The results are shown in Figure 9. We observe that as the half-life of insulin increases, the response of glucagon to low blood glucose levels becomes significantly more impaired, which leads to longer and more severe hypoglycemia. The failure of glucagon to respond to low blood glucose levels in type 1 diabetes is well documented [30]. Our simulations suggest that a main cause of the lack of response of glucagon is due to the clearance of insulin. In the original simulations, we see that insulin peaks and then decreases rapidly allowing for glucagon to respond to lower blood glucose levels. However, in the case of exogenous insulin with longer half-lives, the reduction in insulin clearance means that glucagon is suppressed for longer and therefore cannot respond appropriately to low blood glucose levels as demonstrated in Figure 9. This also suggests that blood glucose levels may be more challenging to regulate in patients administering insulin, since if glucagon is not responding appropriately, a patient may no longer rely on the inhibition of glucagon to regulate glucose levels. This is especially clear in Figure 4, where we see that when glucose undershoots its steady state value, glucagon is able to quickly reduce levels back to steady state. This demonstrates the degree to which the body relies on the anti-insulin properties of glucagon to regulate glucose levels, and highlights the importance of developing insulins that have half-lives closely resembling endogenous insulin in order to avoid impairing the function of glucagon.

## Conclusions and Future Work

In this work we developed a three-compartment model of insulin, glucose and glucagon dynamics and validated our model by fitting it to the median of publicly available IVGTT pig data. We demonstrated that a two-compartment model of insulin and glucose was not sufficient to model the IVGTT pig data (see Supplementary Material Section 7.2), and thus highlighted the importance of glucagon in the insulin-glucose regulatory system. We then removed the feedback of glucose on insulin production (see Figure 7), which allowed us to investigate the impact the loss of communication between insulin and the system has on glucose and, perhaps more interestingly, glucagon dynamics. This is a simplified phenomena of what occurs in both type 1 and type 2 diabetes.

Overall, our model highlights the importance of glucagon in insulin-glucose dynamics. Without glucagon, we were unable to capture insulin-glucose dynamics well (Supplemental Material Figure 11). Furthermore, when glucose feedback on insulin production was removed, changes in insulin from the endogenous response leads to impairment of glucagon response (Figures 8 and 9); inhibition of glucagon subsequently leads to prolonged and severe hypoglycemia (Figures 8b and 9). Moreover, Figure 4 demon-strates the importance of the synergy between insulin and glucagon to optimally regulate glucose. Specifically, we observe that an undershoot of the glucose steady state value is rapidly counteracted via glucagon; note that this is not observed in Figure 9, where feedback was removed. Moreover, upon the removal of feedback, it is still possible for glucagon to regulate the system, but only if the insulin profile is adjusted (see Figure 8(c)); this highlights the important role of communication between glucagon and insulin to ensure that both are responding properly to glucose levels. However, for patients with diabetes, achieving insulin dynamics via external mechanisms similar to endogenous insulin secretion can be nearly impossible, and thus dysfunction of glucagon is often observed.

Future work will continue to focus on the deregulation of insulin. We intend to use the proposed model (1) -(3) to understand how the progression of type 1 diabetes impacts this system by incorporating both the immune system and *β* cell loss. We will then combine this model to the insulin-glucose regulatory system via the insulin compartment (i.e. decreased *β* cell mass implies reduced insulin secretion) to understand how insulin, glucose and glucagon become increasingly deregulated throughout the progression of the disease.

## 7. Supplementary Material

### 7.1. Uniqueness of steady state of insulin-glucose-glucagon model

In this section, we show that system (1) -(3) has a unique positive steady state. Let (*I*_*∗*_,*G*_*∗*_,*G*_*𝓁,∗*_) denote a steady state of system (1) -(3). Thus, (*I*_*∗*_,*G*_*∗*_,*G*_*𝓁,∗*_) satisfy the following system of equations:

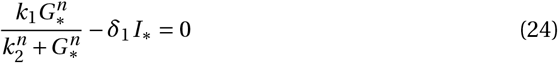

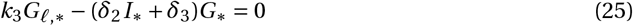

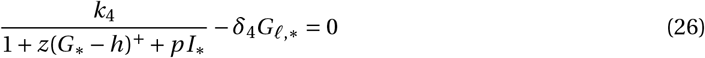

Equation (24) implies that

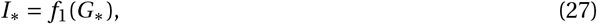

where

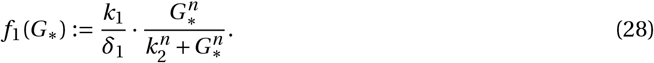

Solving (25) for *G*_*𝓁,∗*_, and using *I*_*∗*_ *= f*_1_(*G*_*∗*_) yields

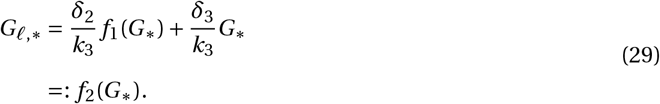

Define

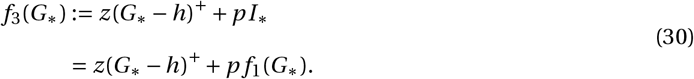

Thus, equation (26) can be written entirely in terms of *G*_*∗*_:

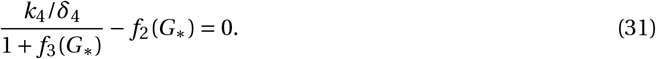

Since *f*_3_(*G*_*∗*_) is increasing and *f*_3_(0) *=* 0, the first term on the left-hand side of (31) is decreasing (as a function of *G*_*∗*_) and takes the value *k*_4_/*δ*_4_ at *G*_*∗*_ *=* 0. The second term, *f*_2_(*G*_*∗*_) is increasing and satisfies *f*_2_(0) *=* 0. Thus, equation (31) has precisely one positive solution *G*_*∗*_. As *G*_*𝓁,∗*_ and *I*_*∗*_ are determined uniquely from *f*_2_ and *f*_1_, respectively, and are also strictly positive (if *G*_*∗*_ is), the claim is shown: system (1) - (3) has a unique positive steady state.

### 7.2. Two Compartment Model

Here we propose a two compartment model of insulin-glucose dynamics; see Section 2 for a discussion of specific expression used in the below system of differential equations.

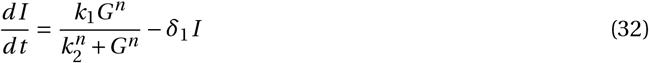

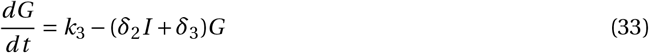

A schematic of the modeled interactions is provided in Figure 10. Note that in system (32) - (33) we do not incorporate the role of glucagon on stimulating glucose production, and thus there is no inhibition of glucose or insulin in this system (see equation (3)).

**Figure 10:**
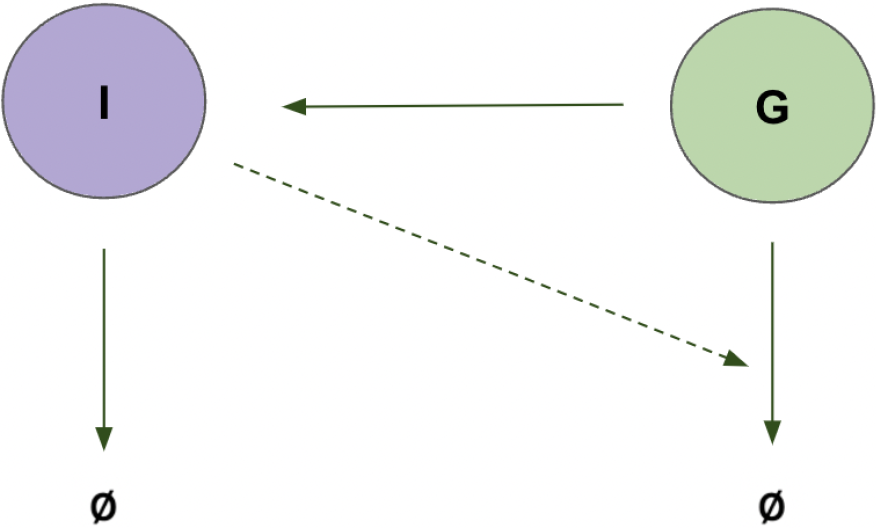
Two compartment insulin-glucose model schematic

**Figure 11:**
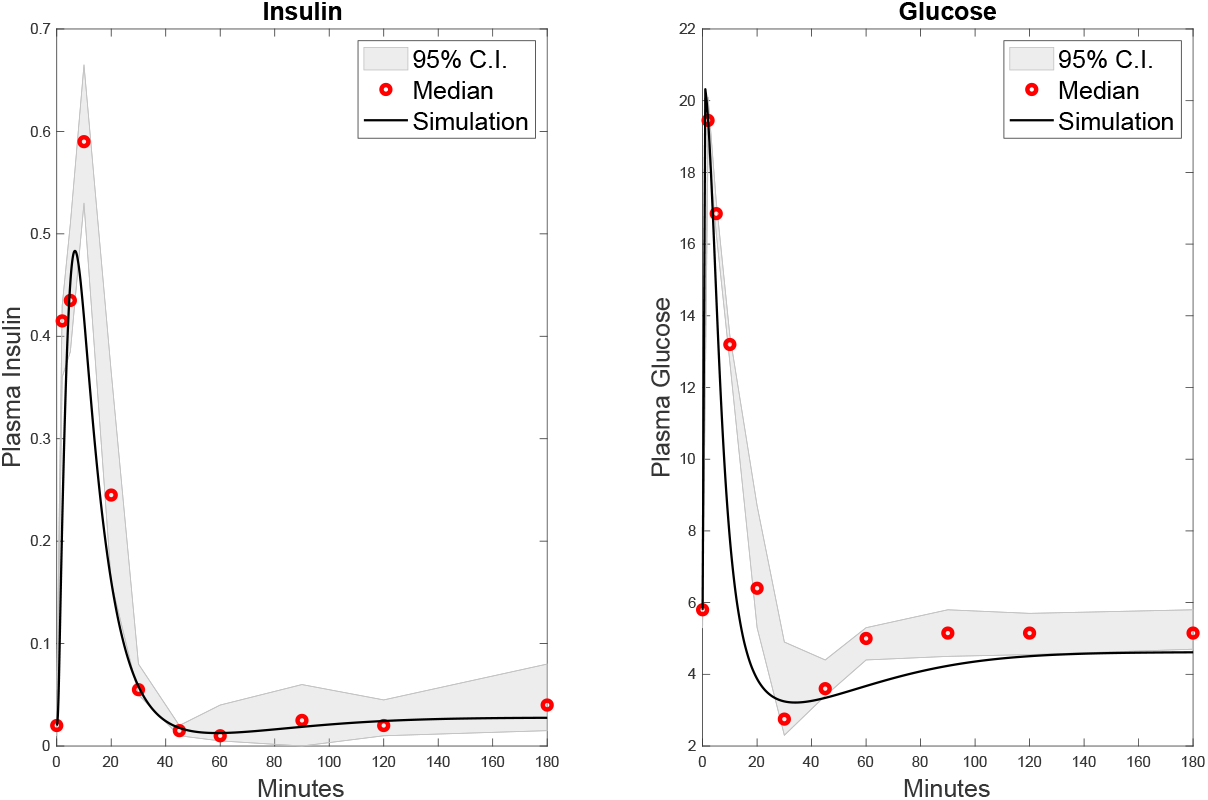
The best fit for the insulin-glucose model 32 - 33. The cost function evaluated at the optimal set of parameters is 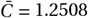 and the adjusted *R*^2^ value is 0.7754. See Table 2 for the corresponding estimated parameters.

The results of fitting the insulin and glucose pig data to system (32) - (33) are provided in Figure 11. Recall that in this case, the cost function that we minimize is given by (11). For details on the external data and calibration methods, see Section 3. Tornado plots to ensure minimality are provided in Figure 12, and the optimal set of parameter values are shown in Table 2.

**Figure 12:**
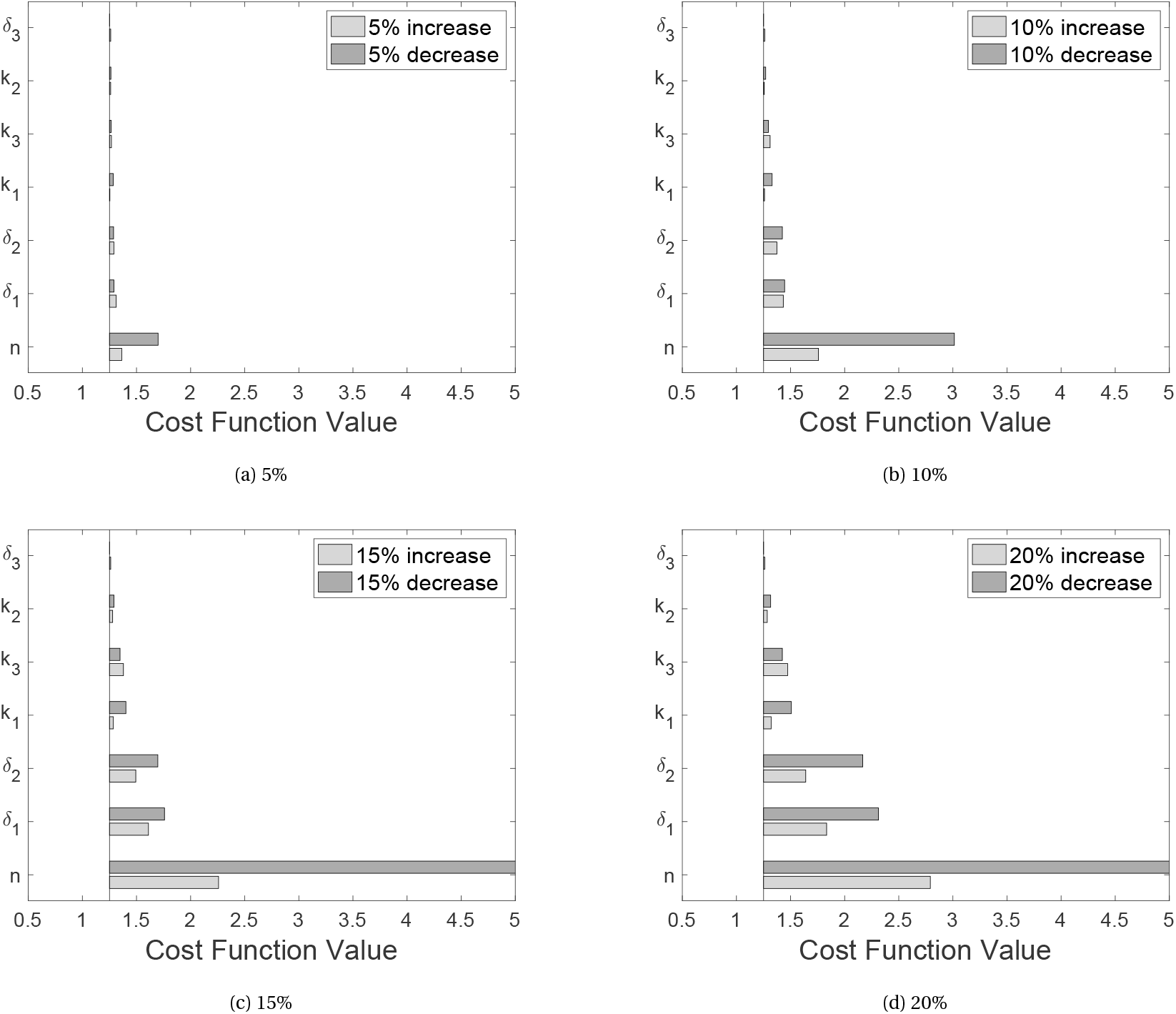
Tornado plots for the change in cost function 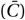 when varying each parameter by the given percentage of its optimal value. The vertical line represents the minimum residual.

### 7.3. Estimating parameters in the minimal model

We provide the best fit solution (Figure 13), estimated parameters (Table 3), and tornado plots (Figure 14) for the minimal model (12) - (14). For details on the external data and calibration methods, see Section 3. A comparison between the insulin-glucose-glucagon model and the minmimal model is provided in Figure 6 in the main text of the article.

**Figure 13:**
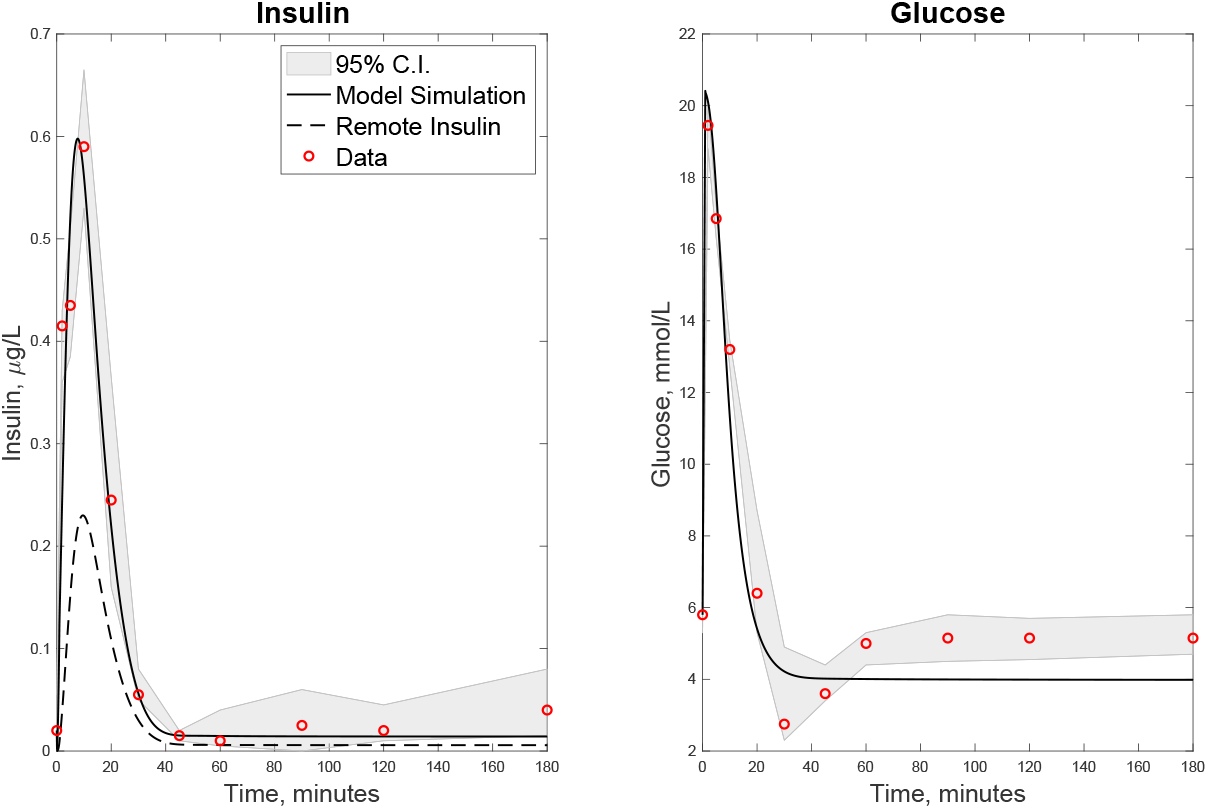
The best fit for the minimal model (12) - (14). The cost function evaluated at the optimal set of parameters is 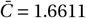. See Table 3 for the corresponding estimated parameters.

**Figure 14:**
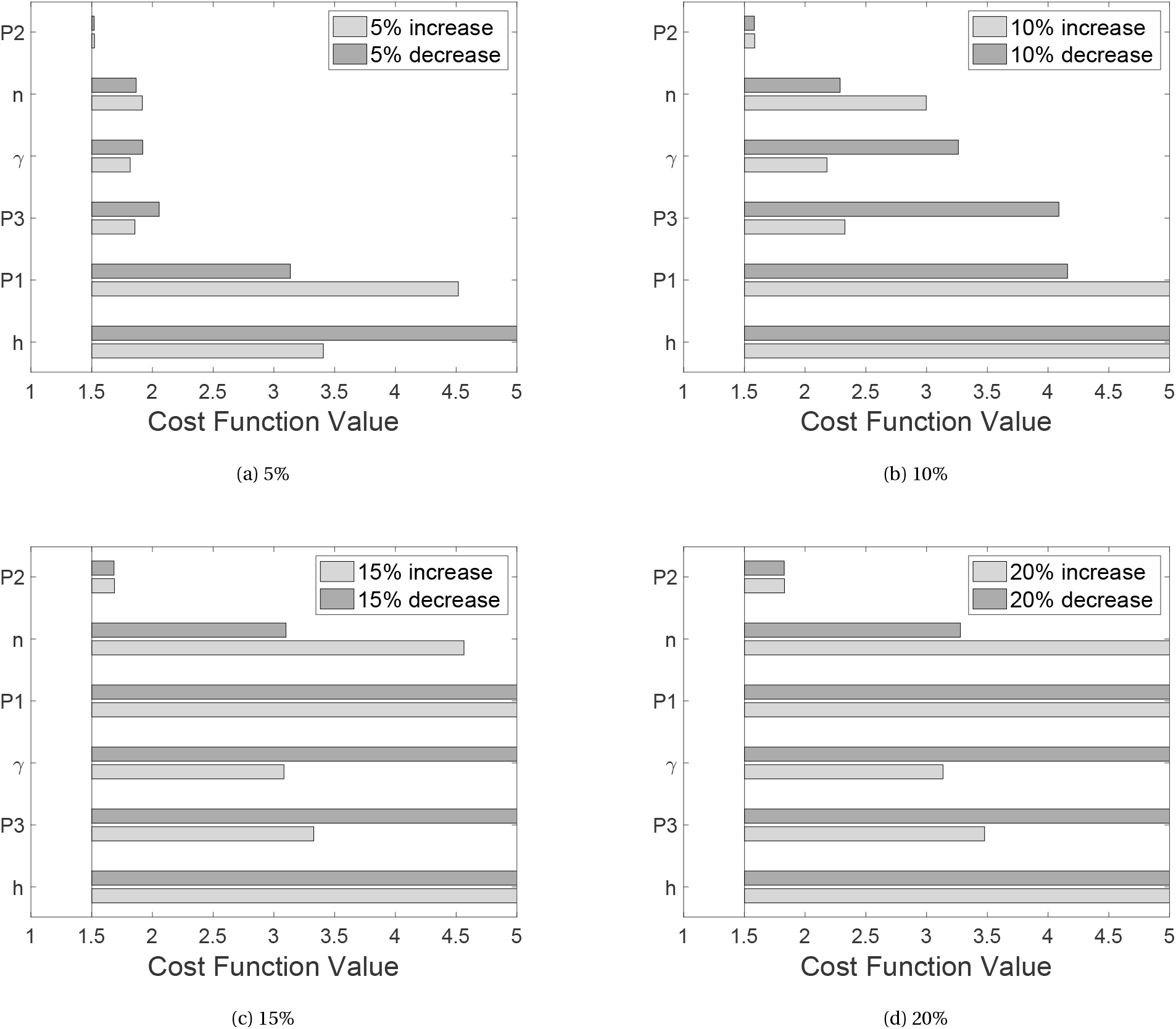
Tornado plots for the change in the cost function 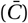 when varying each parameter by the given percentage of its optimal value. The vertical line represents the minimum residual.

